# Multi-Level Attention Graph Neural Network for Clinically Interpretable Pathway-Level Biomarkers Discovery

**DOI:** 10.1101/2020.12.03.409755

**Authors:** Xiaohan Xing, Fan Yang, Hang Li, Jun Zhang, Yu Zhao, Biaobin Jiang, Junzhou Huang, Max Q.-H. Meng, Jianhua Yao

## Abstract

Precision medicine, regarded as the future of healthcare, is gaining increasing attention these years. As an essential part of precision medicine, clinical omics have been successfully applied in disease diagnosis and prognosis using machine learning techniques. However, existing methods mainly make predictions based on gene-level individual features or their random combinations, none of the previous work has considered the activation of signaling pathways. Therefore, the model interpretability and accuracy are limited, and reasonable signaling pathways are yet to be discovered. In this paper, we propose a novel multi-level attention graph neural network (MLA-GNN), which applies weighted correlation network analysis (WGCNA) to format the omic data of each patient into graph-structured data, and then constructs multi-level graph features, and fuses them through a well-designed multi-level graph feature fully fusion (MGFFF) module to conduct multi-task prediction. Moreover, a novel full-gradient graph saliency mechanism is developed to make the MLA-GNN interpretable. MLA-GNN achieves state-of-the-art performance on transcriptomic data from TCGA-LGG/TCGA-GBM and proteomic data from COVID-19/non-COVID-19 patient sera. More importantly, the proposed model’s decision can be interpreted in the signaling pathway level and is consistent with the clinical understanding.

## Introduction

Precision medicine aims at enhancing the treatment effect compared to one-fits-all products by providing personalized treatment plans, medical decisions, and products (Yau 2019). One natural way to assign patients into customized subgroups is through clinical omic data, such as the data from genomics, transcriptomics, and proteomics. Among the omic data, biomarkers which indicate the biological state or disease progression are generally applied for disease diagnosis and prognosis. Researchers usually detect gene mutations or specific individual gene expression profiles (GEPs) to discover biomarkers. The GEPs can be represented either at the transcription level using transcriptomics or at the protein level using proteomics. In addition to the above biomarkers, pathway-level biomarkers have recently been proposed and proven to have unique advantages in medical outcome predictions (Ben-Hamo et al. 2020).

Although omic data have been studied in disease diagnosis and prognosis (Zhang et al. 2018; Shen et al. 2020; Chen et al. 2017), there remain some problems to be solved. First, existing methods mainly make predictions based on individual GEPs or their random combinations, which cannot correctly reflect the complex disease mechanism, thus yielding limited performance. Second, relying on random combinations of GEPs, the methods have poor generalizability due to the batch effects (Haghverdi et al. 2018), which are caused by the instability of GEPs on different batches of data due to the differences in experiment times, handlers, reagent lots, etc. Third, the biological regulatory network is a cascading amplification process. A small change in the transcription factor can amplify its signal, and result in a great change of functional proteins. Statistical analysis can only discover functional proteins with great changes while ignoring the driven factor of the disease, which are clinically relevant and usually used as the drug target. Furthermore, current research mainly focuses on transcriptomics while ignoring the irreplaceability of proteomics. In fact, some disease development (e.g., the infectious disease COVID-19) can only be reflected and detected by proteomics (e.g., sera GEPs). Without enough proteomics researches, current methods cannot comprehensively reveal the underlying disease mechanism.

To solve these problems, we thoroughly inspect the omic data and delicately design our method. The omic data is non-Euclidean since the GEPs do not form grid-structures like image data, meaning the convolutional neural networks (CNNs) that achieve great success in computer vision are not suitable for processing the omic data. Instead, the omic data can be represented in a graph structure since the transcription factors and the functional proteins make up a hierarchical regulatory network in biology. Considering these characteristics, we employ graph neural networks (GNNs) to exploit the information contained in the non-Euclidean graph-structured omic data. Specifically, we format the omic data into WGCNA graphs, which connect the GEPs that perform similar functions or on the same signaling pathway, to mimic the GEP connections in the organism. Then, we propose a multi-level attention graph neural network (MLA-GNN) to imitate the hierarchical regulatory network. As the layer deepens, the MLA-GNN gradually integrates the GEPs on the same signaling pathway, and amplify the contribution of the driven factors, thus can better reveal the underlying disease mechanism or biological process. Furthermore, we develop a full-gradient graph saliency mechanism to interpret the model performance and discover pathway-level biomarkers. By imitating the biological regulatory process and identifying pathway-level biomarkers, the proposed MLA-GNN is robust to batch effects, resulting in efficiency and accuracy on prediction tasks on the transcriptomic data as well as the proteomic data. Our contributions in this work can be summarized as follows:

- We propose a concise and robust MLA-GNN on omic data to imitate biological processes and discover pathway-level biomarkers, supporting survival prediction and classification. To the best of our knowledge, MLA-GNN is the first work that utilizes GNNs to explore the prior structured information contained in the WGCNA graphs.
- We develop a novel full-gradient graph saliency (FGS) mechanism for the interpretation of GNN-based models and validate its superiority on the proposed MLA-GNN.
- Using the FGS mechanism, the MLA-GNN can discover clinically interpretable pathway-level biomarkers, which are relevant but cannot be discovered by existing methods.
- We perform extensive experiments on different tasks using transcriptomic data and proteomic data. The results demonstrate that the superiority and robustness of the MLA-GNN compared to state-of-the-art methods.

## Related Work

In this section, we introduce the existing methods for clinical omics analysis.

### Statistical Approaches for GEP Analysis

At present, most of the clinical analysis on GEPs are based on statistical approaches (Chen et al. 2017; Zhang et al. 2018; Yang et al. 2018), where differentially expressed proteins are calculated using a certain fold change and p-value threshold by t-test or its variants. Unsupervised learning methods, especially hierarchical clustering, are often used for disease subtyping. For instance, comprehensive LUAD proteogenomics exposes multi-omic clusters and immune subtypes (Gillette et al. 2020). However, it cannot provide a specific decision bound to predict the efficacy or the subtype of each patient, and often has a high false discovery rate due to the “large p, small n” problem (Diao and Vidyashankar 2013), hindering its clinical application.

### Machine Learning-based GEP Analysis Methods

In recent years, machine learning has been widely employed in the medical field (Zhao et al. 2020; Wang et al. 2020; Liang et al. 2020). Encouraged by these studies, an increasing number of machine learning-based methods are applied to omic data. Random forest is reported to successfully predict the risk of preterm delivery of pregnant women based on cell-free RNA (Ngo et al. 2018). Support Vector Machine (SVM) using single-cell transcriptomic data is applied to predict brain development through distinguishing neocortical cells and neural progenitor cells (Hu et al. 2016). However, these methods require great efforts in hand-crafted feature engineering and have limited generalizability.

### Deep Learning-based GEP Analysis Methods

Deep learning, as the most recent iteration of the machine learning method, has been applied to GEP analysis. Self-Normalizing Network (SNN), as a variant of fully connected neural networks, achieves state-of-the-art performance on cancer diagnosis and prognosis tasks using RNAseq data (Chen et al. 2019). However, the features in SNN are randomly combined based on the weight matrix between adjacent layers, such random combination cannot fully exploit the inherent structure information of the omic data. Recently, GNNs (Kipf and Welling 2016; Veličković et al. 2017) have been applied to predict the node-level embeddings of Protein-Protein Interaction (PPI) graphs. However, graph-level predictions, which have great clinical significance in predicting the phenotype or survival time of each patient, are under-studied.

## Methods

The overview of the proposed MLA-GNN is illustrated in Figure 1. Given the training data *X*^*N*×*K*^ for *N* patients (*K* GEPs for each patient), we first construct the edge matrix *E*^*K*×*K*^ through WGCNA analysis (Langfelder and Horvath 2008). Then, each patient can be represented by a graph *G*_1_ = *G*(*V*^*K*×1^, *E*^*K*×*K*^), where *V*^*K*×1^ represents the features of *K* nodes and the edge matrix *E*^*K*×*K*^ denotes the edge connections in the graph. We then utilize several graph attention (GAT) layers (Veličković et al. 2017) to construct hierarchical graph features *G*_2_ and *G*_3_ from the WGCNA graph *G*_1_. In the proposed multi-level graph feature fully fusion (MGFFF) module, the multi-level graph features are fused after linear projection (LP) and vectorization. Then, the fused feature is sent to the last stage of the pipeline, a sequential network for multi-task prediction, such as disease classification and survival prediction. Furthermore, we propose a novel full-gradient graph saliency (FGS) mechanism to reveal the importance of each node on the graph, thus providing clinical interpretation for the proposed model.

**Figure 1:**
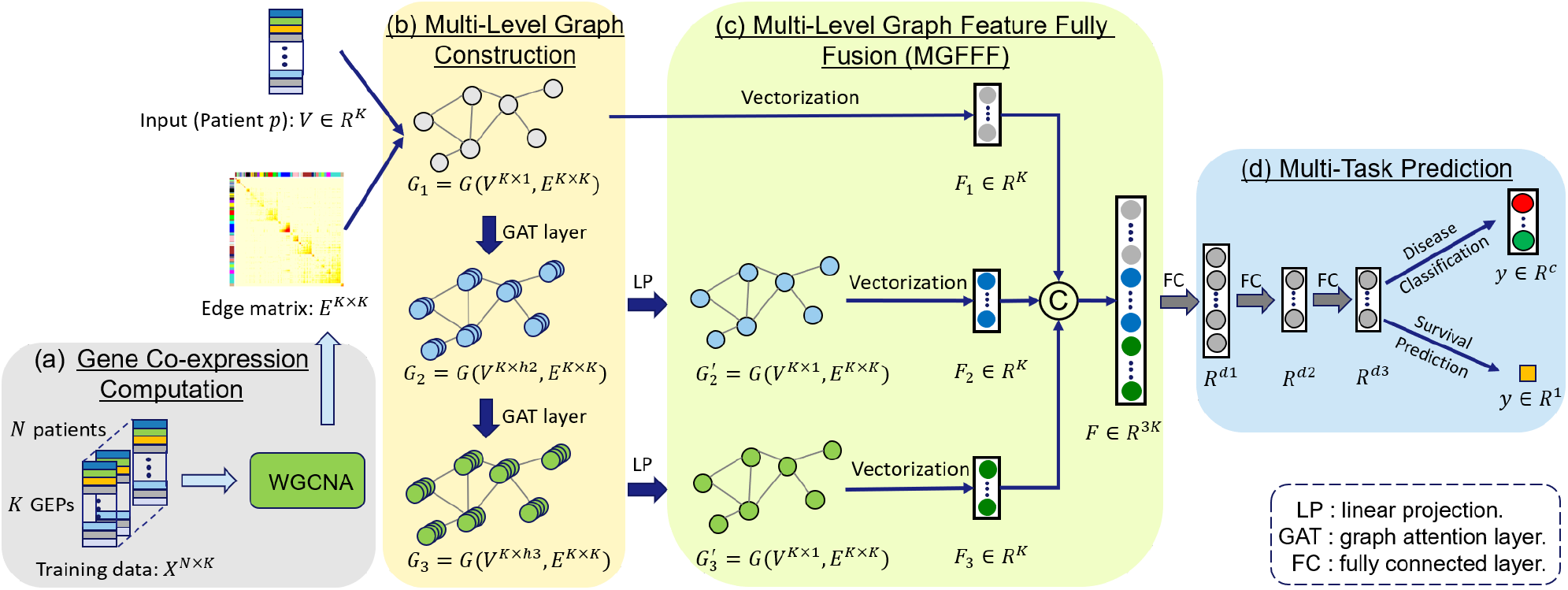
Overview of the proposed Multi-Level Attention Graph Neural Network (MLA-GNN). (a) Gene Co-expression Computation module performs weighted correlation network analysis (WGCNA) on the training data to produce the edge matrix. (b) Multi-Level Graph Construction module builds multi-level graphs through GAT layers. (c) Multi-Level Graph Feature Fully Fusion (MGFFF) module integrates local gene-level features and global pathway-level features. (d) Multi-Task Prediction module conducts multiple medical tasks, such as disease classification and survival prediction.

### Gene Co-expression Computation

In order to employ GNNs for omics analysis, the first step is to format the omic data of each patient into a graph which is specified by an input feature *V* and an edge matrix *E*. Suppose each patient is represented by *K* GEPs, the feature can be represented by *V*^*K*×1^, which corresponds to a graph with *K* nodes, with each node contains the expression of a gene. As shown in Figure 1 (a), we perform gene co-expression computation through WGCNA analysis to calculate the edge matrix *E*. Specifically, for the training data *X*^*N*×*K*^ (where *N* denotes the number of patients in the training set), the expression profile of each gene (node) is characterized by an *N*-dimension vector. For any two nodes *v_i_* and *v_j_* ∈ *R^N^*, their pairwise correlation *A_ij_* is calculated as

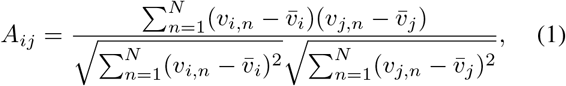

where 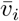 and 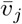 are the average features of the node *v_i_* and *v_j_*. In this way, the nodes with similar gene expressions are connected with larger adjacency values.

To construct the edge matrix *E*, we binarize the continuous values in the adjacency matrix *A* through

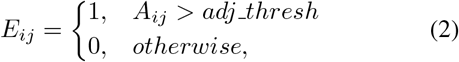

where the hyper parameter *adj_thresh* is optimized by an automated machine learning (Waldrop, Youn, and Patterson 2014) algorithm. Note that the edge matrix *E* is calculated based on all training data, thus is not patient-specific and does not need to be computed repeatedly.

Utilizing the edge matrix *E*, we format each patient into a WGCNA graph *G*_1_ = *G*(*V*^*K*×1^, *E*^*K*×*K*^). In the WGCNA graph, nodes with similar gene expressions are connected by edges, while others are not. It is reported in (Langfelder and Horvath 2008) that genes with similar expressions usually conduct similar functions and are more likely to be mapped to the same signaling pathway. Therefore, the genes on a signaling pathway are naturally connected in the WGCNA graph, thus could be processed to extract pathway-level information that cannot be discovered by existing methods.

### Multi-Level Graph Construction

For each patient, the constructed WGCNA graph *G*_1_ = *G*(*V*^*K*×1^, *E*^*K*×*K*^) is fed into a stack of GAT layers to construct multi-level graph features. GAT layer is an advanced graph convolutional layer which outputs each node feature as the weighted combination of its neighboring nodes and the node itself. A self-attention mechanism is performed to compute the attention coefficient between the node and its neighbors. For example, the attention coefficient between the node *v_i_* ∈ *R^h^* and its neighbor *v_j_* ∈ *R^h^* is computed as

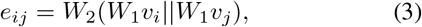

where *W*_1_ is the weight parameter of a fully connected layer, || denotes the concatenation operation, and a fully connected layer with weight parameter *W*_2_ ∈ *R*^2*h*^ encodes the correlation between the node *i* and node *j*. Then, the attention coefficients are normalized using the softmax function

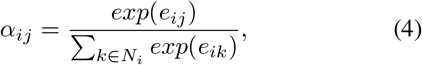

there *k* ∈ *N_i_* denotes all first-order neighbors of the node *i* and the node itself. After that, the normalized attention coefficients *α_ij_* are used to compute a linear combination of the corresponding features as the output features for the node *i*:

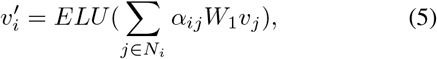

where ELU, a combination of Sigmoid and ReLU, is the non-linear activation function. According to Eq. 5, the features of neighboring nodes with high similarity are integrated into the target node with large weights.

As aforementioned, GEPs on the same signaling pathway are more likely to be connected in the WGCNA graphs. Therefore, the output feature of each node in the graph after the GAT layer is the weighted combination of the GEP features on the signaling pathway, which is a natural way to extract the feature representations in the pathway-level. To the best of our knowledge, this work is the first to utilize GNNs to explicitly explore the prior structured information contained in the WGCNA graphs.

### Multi-Level Graph Feature Fully Fusion

As shown in Figure 1, there are three different levels of graphs, including the input graph *G*_1_, and the generated high-level graphs *G*_2_ and *G*_3_. These graphs, although having the same number of nodes, contain hierarchical information. Specifically, the feature of a node in *G*_1_ represents the expression of a specific gene, while each node in *G*_2_ or *G*_3_ contains features from many GEPs in a signaling pathway. Therefore, we denote the features from *G*_1_ as local GEP features and those from *G*_2_ and *G*_3_ as global pathway features.

Since both local GEP features and global pathway features are important in omics representation learning and phenotype prediction (Ben-Hamo et al. 2020), we fuse the multi-level graph features to produce more discriminative feature representation. However, due to the different node dimensions in multi-level features, direct concatenation would put more weights on the graph at a higher level. To this end, we first perform linear projection by a fully connected layer to generate high-level graph features 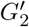 and 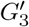 with reduced node dimension. Then, the features *G*_1_, 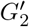, and 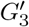 are vectorized to generate three same-dimension feature vectors *F*_1_, *F*_2_, *F*_3_ ∈ *R^K^*, which are concatenated to produce the fused feature *F* ∈ *R*^3*K*^. By this design, the network could adaptively select the most meaningful information during the training process and find the desired representation for each node, which will lead to a leap in model capacity. Benefiting from the global pathway-level information, the fused feature *F* ∈ *R*^3*K*^ enables the model to discover high-level biomarkers (e.g., transcription factors) that cannot be discovered by the original input feature *F*_1_ ∈ *R^K^*. This is of great significance to bioinformatics research and clinical applications.

It is noteworthy that different from the common methods which generate graph representations by pooling across all the nodes, our method compresses the features inside each node while keeping the node structure in the graph. This is a specific design in this work, and the reason is that each node corresponding to an individual GEP has its special biological meaning, thus cannot be compressed across nodes.

### Multi-Task Prediction

The proposed MLA-GNN is designed to address multiple different clinical tasks. As shown in Figure 1, the fused feature *F* is encoded by a sequential fully connected layers to perform disease classification and survival prediction in the multi-task prediction module.

For the disease classification task, the output *y* ∈ *R^c^* denotes the probability scores of *c* classes. This task is optimized by the cross-entropy loss. For the survival prediction task, the output *y* ∈ *R*^1^ denotes “hazard ratio”. And the cox loss is calculated as

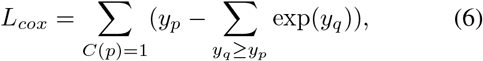

where *C*(*p*) = 1 means the *p*-th patient is censored, and only censored patients are included in the cox loss computation.

Note that the proposed model degrades to SNN (Chen et al. 2019) if we utilize the input feature *F*_1_ rather than the fused feature *F* for multi-task prediction.

### Full-Gradient Graph Saliency

Currently, increasing concern regarding the interpretability of deep neural networks has been raised. A clinically interpretable model, which can reveal the working mechanism and improve the credibility of the model, is especially demanded in the clinical community. In this paper, considering the success of full-gradient saliency in CNNs (Srinivas and Fleuret 2019), we develop a novel full-gradient graph saliency (FGS) mechanism to interpret GNNs and provide clinical explanations for the proposed MLA-GNN.

In the MLA-GNN, the node on the input graph *G*_1_ represents the expression of an individual gene and may have limited importance (local importance). However, the node on *G*_2_ and *G*_3_ combining a group of GEPs may be critical if they form an important signaling pathway (global importance). From the view of the clinical community, the local importance could reveal low-level functional biomarkers, and the global importance help discovers high-level regulatory factors. Thus, they are both crucial in clinical interpretation and biological research. To this end, the FGS mechanism is designed to reveal node importance by integrating the gradients of multi-level graph features.

First, for the *p*-th patient, we deduce level-wise feature importance by computing the gradients of target *t_p_* over the input graph *G*_1_ and over the intermediate graphs 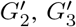. Specifically, for *l* ∈ {1, 2, 3} corresponding to the graph features of different levels (i.e., *G*_1_, 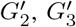), the gradient for the *i*-th node of the *p*-th patient is calculated by

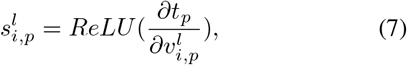

where 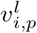 is the feature of the *i*-th node in the *l*-th level of the *p*-th patient, ReLU is the activation function which is utilized to remove negative gradient response. According to the different tasks, the target *t_p_* in Eq. 7 is defined as

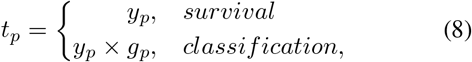

where *y_p_* and *g_p_* are the prediction and ground-truth labels of the *p*-th patient.

Then, we compute the saliency score *s_i_* for the *i*-th node by aggregating the gradients cross three levels and all patients, which is defined as

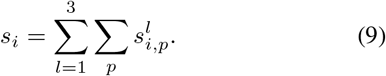

The FGS mechanism would reveal the contribution of each GEP to the final prediction. The top GEPs with high saliency scores are recognized as biomarkers and will be analyzed by biological methods for further biological insights.

## Experiment

To comprehensively evaluate the performance of MLA-GNN, we conduct experiments on two public datasets, including transcriptomic data and proteomic data separately, for the tasks of survival prediction, histological grading, and COVID-19 diagnosis. Moreover, taking the TCGA-LGG/TCGA-GBM dataset as an example, we show the paradigm of the model interpretability which is an important breakthrough in clinical omics.

### Datasets

The summary of the datasets used in the experiments is shown in Table 1.

**Table 1:** Data summary

| Dataset | Data source | # Patients | # GEPs | Type of omics | Tasks |
| --- | --- | --- | --- | --- | --- |
| Glioma | TCGA-LGG & TCGA-GBM | 769 | 240 | transcriptomics | survival outcome & histological grading |
| COVID-19 | COVID-19 patients sera | 70 | 791 | proteomics | COVID-19 diagnosis |

#### Glioma Dataset: RNAseq of glioma patients for survival prediction and histological grading

The glioma cases are collected from the TCGA-GBM and TCGA-LGG projects. The dataset contains a total of 769 patients. Each case contains 240-dimensional RNAseq data curated from the TCGA and cBioprotal platforms (Cerami et al. 2012). The corresponding clinical information includes survival outcomes and histological grading (grade II, grade III, and grade IV), as the labels to be predicted by the models.

#### COVID-19 Dataset: Proteomic data of the COVID-19 patients sera for diagnosis

The outbreak of the COVID-19 pandemic has brought a global crisis. Recently, the sera proteomic data from some COVID-19 cases have been released (Shen et al. 2020). We also apply MLA-GNN to the classification of COVID-19 patients, contributing to the understanding and auxiliary diagnosis of COVID-19. The dataset contains 34 COVID-19 patients and 36 non-COVID-19 patients, with 791 proteins identified in the sera samples.

### Implementation Details

We implement the proposed MLA-GNN with Pytorch (Paszke et al. 2017) and Pytorch Geometric library (Fey and Lenssen 2019). The model is trained in an end-to-end manner utilizing Adam optimizer and with batch size set to 8.

The learning rate is initialized as 0.002 and linearly decayed in the training process. For the COVID-19 dataset, we employ the SelectKBest function in the scikit-learn package to reduce the feature dimensions and use the selected features as input to the proposed model. To tackle the class imbalance problem, we employ the “WeightedRandomSampler” strategy to prepare each training batch data.

To validate the effectiveness and robustness of the proposed method, we conduct cross-validation in the experiments. Specifically, 15-fold cross-validation is performed on the glioma dataset, to be consistent with (Chen et al. 2019) for a fair comparison. For the COVID-19 dataset, we perform 5-fold cross-validation considering the relatively small amount of data.

### Evaluation

For the survival prediction task, performance of the MLA-GNN is compared with the SNN (Chen et al. 2019), cox-PH model (Cox 1972), and cox-nnet (Ching and Garmire 2018). The performances of these models are evaluated with c-index, which is generally employed to measure the performance of survival prediction.

For the histological grading and COVID-19 diagnosis tasks, the performance of the MLA-GNN is compared with the SNN (Chen et al. 2019), SVM (Suykens and Vandewalle 1999), and Random Forest (Liaw, Wiener et al. 2002). These models are evaluated by overall accuracy, precision, recall, and F1-score.

To evaluate the effectiveness of feature fusion in the MGFFF module, we conduct ablation studies to compare the performance of the fused feature *F* and the single-level feature *F*_1_, *F*_2_, and *F*_3_. For clarity, we denote the model with a single-level feature as Single-Level Attention Graph Neural Network (SLA-GNN). Therefore, the models with feature *F*_1_, *F*_2_, and *F*_3_ are named as SLA-GNN-level1 (which is the same as the SNN model), SLA-GNN-level2, and SLA-GNN-level3, respectively.

## Results and Discussion

### Experimental Results

#### Performance on the glioma dataset for survival prediction and histological grading

Table 2 shows the model performance on the survival prediction task of the glioma dataset. The MLA-GNN achieves the c-index of 0.7620, outperforming the cox-PH, cox-nnet, and the current state-of-the-art SNN by a large margin. This result validates the superiority of the proposed MLA-GNN. Furthermore, we can observe performance gains of the MLA-GNN over SLA-GNNs at different levels. This indicates that the fused feature *F* (used in MLA-GNN) is more discriminative than the single-level features (used in SLA-GNNs), thus demonstrating the effectiveness of feature fusion in the MGFFF module.

**Table 2:** Model performance on the survival prediction task of the glioma dataset

| Model | c-index |
| --- | --- |
| cox-PH (Cox 1972) | $0.6911 \pm 0.0748$ |
| cox-nnet (Ching and Garmire 2018) | $0.7152 \pm 0.0730$ |
| SNN* (Chen et al. 2019) | $0.7286 \pm 0.0744$ |
| MLA-GNN (Ours) | **$0.7620 \pm 0.0682$** |
| SLA-GNN-level2 | $0.7382 \pm 0.0674$ |
| SLA-GNN-level3 | $0.7266 \pm 0.0723$ |

\* The SNN can also be called as SLA-GNN-level1.

For the histological grading task, we present the model performance in Table 3. Our proposed MLA-GNN achieves an accuracy of 0.6920, significantly outperforming two commonly used machine learning methods (i.e., SVM and Random Forest), and the deep neural network SNN. Furthermore, compared with SLA-GNNs which make predictions based on single-level features, the MLA-GNN achieves better performance since more comprehensive information is represented by the multi-level fused feature *F*.

**Table 3:**
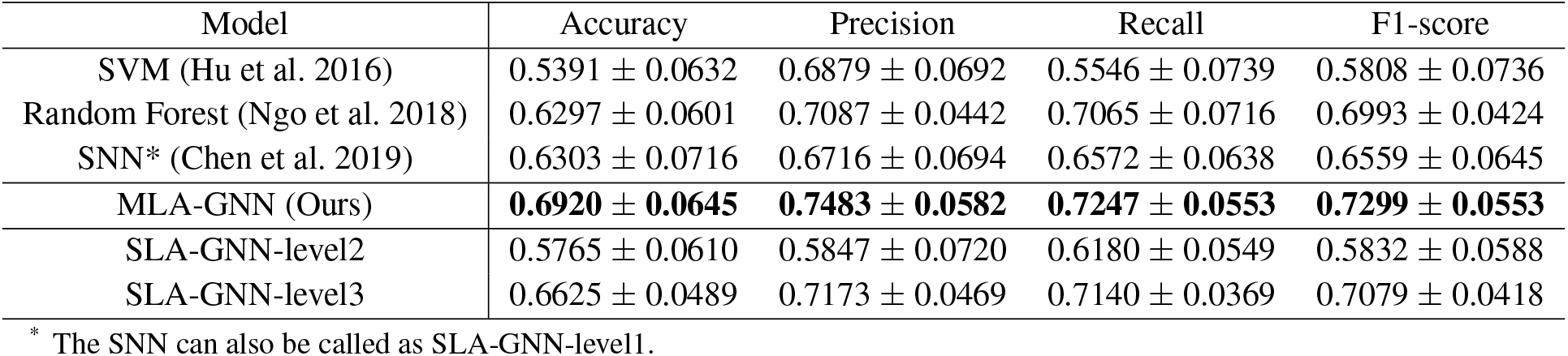
Model performance on the histological grading task of glioma dataset

| Model | Accuracy | Precision | Recall | F1-score |
| --- | --- | --- | --- | --- |
| SVM (Hu et al. 2016) | $0.5391 \pm 0.0632$ | $0.6879 \pm 0.0692$ | $0.5546 \pm 0.0739$ | $0.5808 \pm 0.0736$ |
| Random Forest (Ngo et al. 2018) | $0.6297 \pm 0.0601$ | $0.7087 \pm 0.0442$ | $0.7065 \pm 0.0716$ | $0.6993 \pm 0.0424$ |
| SNN* (Chen et al. 2019) | $0.6303 \pm 0.0716$ | $0.6716 \pm 0.0694$ | $0.6572 \pm 0.0638$ | $0.6559 \pm 0.0645$ |
| MLA-GNN (Ours) | **$0.6920 \pm 0.0645$** | **$0.7483 \pm 0.0582$** | **$0.7247 \pm 0.0553$** | **$0.7299 \pm 0.0553$** |
| SLA-GNN-level2 | $0.5765 \pm 0.0610$ | $0.5847 \pm 0.0720$ | $0.6180 \pm 0.0549$ | $0.5832 \pm 0.0588$ |
| SLA-GNN-level3 | $0.6625 \pm 0.0489$ | $0.7173 \pm 0.0469$ | $0.7140 \pm 0.0369$ | $0.7079 \pm 0.0418$ |

\* The SNN can also be called as SLA-GNN-level1.

#### Performance on the COVID-19 dataset for diagnosis

Comparing with the histological grading of the glioma dataset, the binary classification of the COVID-19 dataset is an easier task since the features of COVID-19 patients and non-COVID-19 patients are very different. The model performance is shown in Table 4. Compared with existing methods (i.e., SVM, Random Forest, and SNN) and SLA-GNNs, our proposed MLA-GNN achieves state-of-the-art performance with an accuracy of 0.9305, which further validates the effectiveness of the MLA-GNN and the MGFFF module on a different task and a different omic dataset.

**Table 4:**
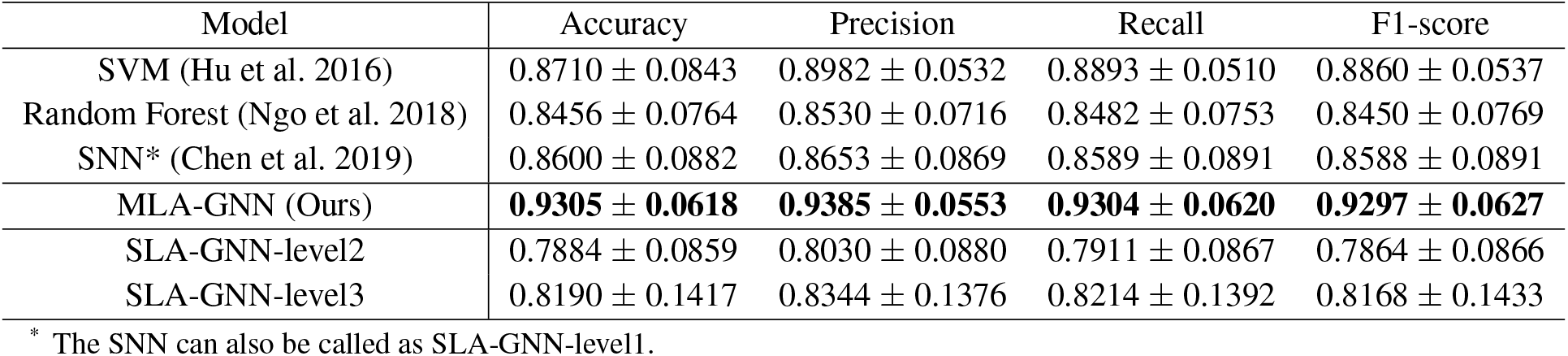
Model performance on the diagnosis of the COVID-19 dataset.

| Model | Accuracy | Precision | Recall | F1-score |
| --- | --- | --- | --- | --- |
| SVM (Hu et al. 2016) | $0.8710 \pm 0.0843$ | $0.8982 \pm 0.0532$ | $0.8893 \pm 0.0510$ | $0.8860 \pm 0.0537$ |
| Random Forest (Ngo et al. 2018) | $0.8456 \pm 0.0764$ | $0.8530 \pm 0.0716$ | $0.8482 \pm 0.0753$ | $0.8450 \pm 0.0769$ |
| SNN* (Chen et al. 2019) | $0.8600 \pm 0.0882$ | $0.8653 \pm 0.0869$ | $0.8589 \pm 0.0891$ | $0.8588 \pm 0.0891$ |
| MLA-GNN (Ours) | **$0.9305 \pm 0.0618$** | **$0.9385 \pm 0.0553$** | **$0.9304 \pm 0.0620$** | **$0.9297 \pm 0.0627$** |
| SLA-GNN-level2 | $0.7884 \pm 0.0859$ | $0.8030 \pm 0.0880$ | $0.7911 \pm 0.0867$ | $0.7864 \pm 0.0866$ |
| SLA-GNN-level3 | $0.8190 \pm 0.1417$ | $0.8344 \pm 0.1376$ | $0.8214 \pm 0.1392$ | $0.8168 \pm 0.1433$ |

\* The SNN can also be called as SLA-GNN-level1.

These results are of great significance since this is the first work to explore and validate the capability of GNNs on proteomic data. With satisfying performance, the MLA-GNN can serve as a tool to assist COVID-19 diagnosis and disease mechanism discovery.

### Discussion

As aforementioned, MLA-GNN achieves the best performance on three different tasks and two different omic datasets. By comparing the results comprehensively, we can gain several insights:

1. Although deep learning-based methods have shown great promises in medical applications, the deep network SNN did not show obvious improvements comparing with other shallow methods. This suggests that the proper design of deep learning algorithms specific to omic data is required to achieve superior performance.
2. In all experiments, the MLA-GNN consistently out-performs SLA-GNNs, thus suggesting the superiority of the fused feature *F* over single-level features. We further take the grading task of the glioma dataset as an example to visualize the feature distributions. As shown in Figure 2, the fused feature *F* tends to be more separable than single-level features, which is consistent with the quantitative results. We conjecture the underlying reason is: Different levels of features may characterize different aspects of a disease. Specifically, low-level graph feature *F*_1_ describes the expression of individual genes (functional GEPs), while high-level features *F*_2_ and *F*_3_ reflect the expressions of a group of genes on a signaling pathway. The development of a disease is a complex mechanism regulated by millions of genes and proteins (which form signaling pathways), thus it cannot be well represented by gene expressions at a single level. In our method, the fused feature *F* integrates information from both the local gene-level and the global pathway-level thus can better reveal the biological mechanism behind diseases.
3. Among the three SLA-GNN models, SLA-GNN-level2 achieves the best performance on the survival prediction of the glioma dataset, SLA-GNN-level3 performs better on glioma grading, while SLA-GNN-level1 (same as SNN) is more accurate for COVID-19 diagnosis. This indicates that features at different levels may be suitable for different tasks and fused features make our model more generalizable.

**Figure 2:**
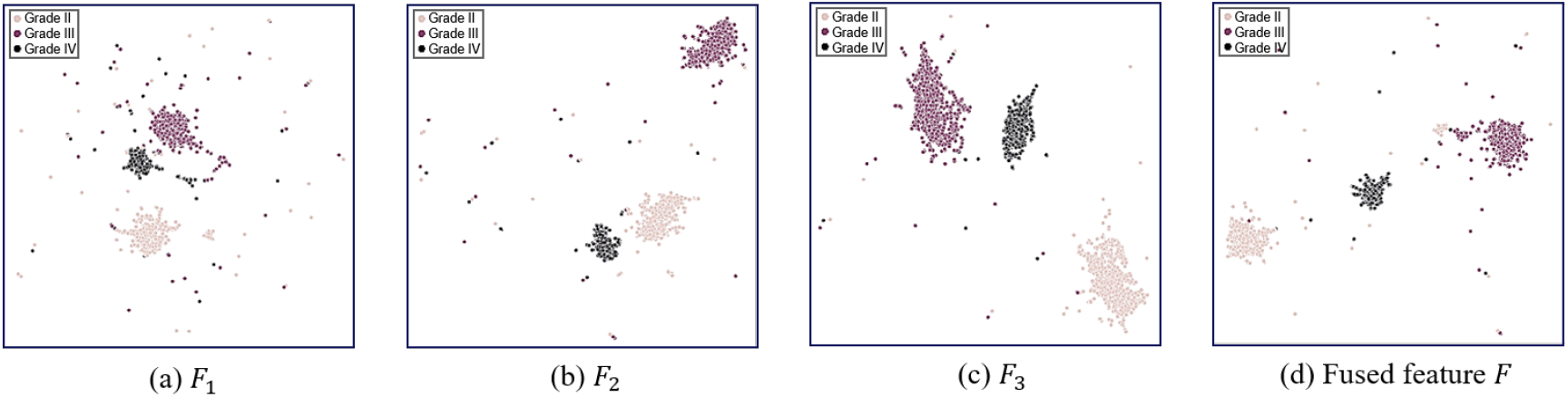
UMAP visualization of the features at different levels for the grading task of the glioma dataset. The fused feature *F* is more separable among different classes (Grade II, Grade III, and Grade IV) than the single-level features *F*_1_, *F*_2_, and *F*_3_.

### Model Interpretability

We use the histological grading task of the glioma dataset as an example to demonstrate the model interpretability.

First, we distinguish the most important nodes utilizing the proposed FGS saliency and perform layer-by-layer analysis. As indicated in Figure 3, the most important node in *F*_1_ is the functional GEP, MKI67, also known as an important biomarker in glioma (Zeng et al. 2015). As the level deepens, the most important node in *F*_3_ becomes the transcription factor, EGFR, which is reported by previous clinical studies as the driven factor for glioma initiation and progression (Huang, Xu, and White 2009). These results show that the existing methods based on the feature *F*_1_ cannot discover the driven factor EGFR. By fusing multi-level graph features through the MGFFF module in the forward propagation and integrating the gradients of multi-level features through the FGS mechanism, our method can discover both the functional GEP (MKI67) and the driven factor (EGFR), thus proving the superiority of our MLA-GNN.

**Figure 3:**
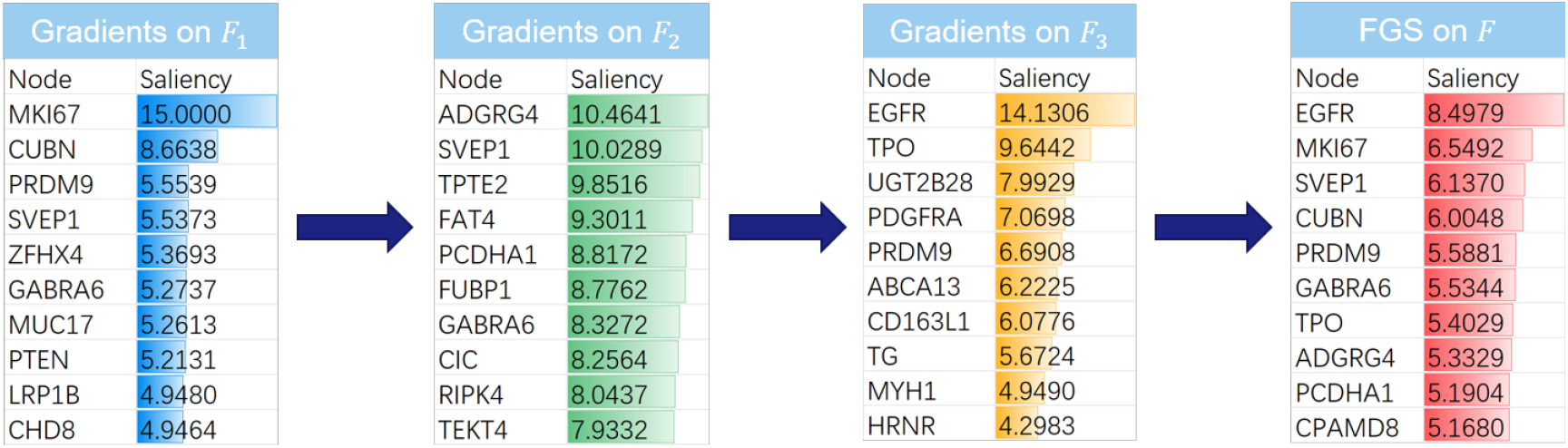
Visualization of the TOP10 important GEPs and their gradient-based saliency scores in each layer and after fusion.

To discover pathway-level biomarkers, we use the Metascape platform (Zhou et al. 2019) to enrich signaling pathways from the most important nodes (GEPs). As shown in Figure 4, the most significant nervous system development and L1CAM interactions are discovered as pathway-level biomarkers. They have been proven to be the most important signaling pathways in glioma progression (Wachowiak et al. 2018). For further verification, we calculate the activation scores of these pathway-level biomarkers in each patient and find that their distributions are significantly different in patients with different histological gradings (i.e., nervous system development: grade II vs grade III p-value 2.21 × 10^−4^, L1CAM interactions: grade II vs grade III p-value 3.33 × 10^−4^). In this way, we prove that the enriched signaling pathways can be used as biomarkers instead of a random combination of individual GEP, throwing light on pathway-level biomarkers discovery using deep learning.

**Figure 4:**
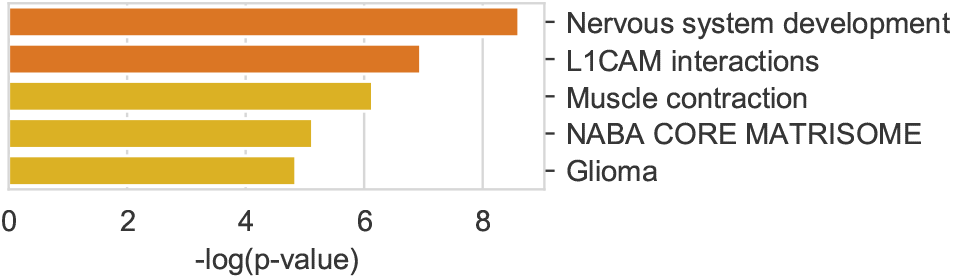
Signaling pathways enriched by the most important GEPs. Larger −*log*(p-value) indicates greater significance in distinguishing different classes.

## Conclusions

In this paper, we propose the pioneer MLA-GNN model on omic data to imitate biological processes and explicitly explore the structured information contained in the WGCNA graphs. On both transcriptomic and proteomic data, extensive experimental results show that MLA-GNN achieves state-of-the-art performance in survival prediction, histological grading, and COVID-19 diagnosis. For model interpretation, we propose a novel full-gradient graph saliency module to distinguish the most important GEPs and discover pathway-level biomarkers. In the future, we will apply our model to more clinical omic data to discover novel pathway-level biomarkers, which will promote the application and interpretation of deep learning in clinical omics. Furthermore, we will explore the application of multi-modality model fusion in precision medicine.

## Notes

### Competing Interest Statement

The authors have declared no competing interest.

## References

Ben-Hamo, R.; Berger, A. J.; Gavert, N.; Miller, M.; Pines, G.; Oren, R.; Pikarsky, E.; Benes, C. H.; Neuman, T.; Zwang, Y.; et al. 2020. Predicting and affecting response to cancer therapy based on pathway-level biomarkers. Nature communications 11(1): 1–16.

Cerami, E.; Gao, J.; Dogrusoz, U.; Gross, B. E.; Sumer, S. O.; Aksoy, B. A.; Jacobsen, A.; Byrne, C. J.; Heuer, M. L.; Larsson, E.; et al. 2012. The cBio cancer genomics portal: an open platform for exploring multidimensional cancer genomics data.

Chen, L.; Yang, F.; Chen, X.; Rao, M.; Zhang, N.-N.; Chen, K.; Deng, H.; Song, J.-P.; and Hu, S.-S. 2017. Comprehensive myocardial proteogenomics profiling reveals C/EBP*α* as the key factor in the lipid storage of ARVC. Journal of Proteome Research 16(8): 2863–2876.

Chen, R. J.; Lu, M. Y.; Wang, J.; Williamson, D. F.; Rodig, S. J.; Lindeman, N. I.; and Mahmood, F. 2019. Pathomic Fusion: An Integrated Framework for Fusing Histopathology and Genomic Features for Cancer Diagnosis and Prognosis. arXiv preprint arXiv:1912.08937.

Ching, T.; and Garmire, L. X. 2018. Cox-nnet: an artificial neural network method for prognosis prediction of high-throughput omics data. PLoS computational biology 14(4): e1006076.

Cox, D. R. 1972. Regression models and life-tables. Journal of the Royal Statistical Society: Series B (Methodological) 34(2): 187–202.

Diao, G.; and Vidyashankar, A. N. 2013. Assessing genome-wide statistical significance for large p small n problems. Genetics 194(3): 781–783.

Fey, M.; and Lenssen, J. E. 2019. Fast graph representation learning with PyTorch Geometric. arXiv preprint arXiv:1903.02428.

Gillette, M. A.; Satpathy, S.; Cao, S.; Dhanasekaran, S. M.; Vasaikar, S. V.; Krug, K.; Petralia, F.; Li, Y.; Liang, W.-W.; Reva, B.; et al. 2020. Proteogenomic characterization reveals therapeutic vulnerabilities in lung adenocarcinoma. Cell 182(1): 200–225.

Haghverdi, L.; Lun, A. T.; Morgan, M. D.; and Marioni, J. C. 2018. Batch effects in single-cell RNA-sequencing data are corrected by matching mutual nearest neighbors. Nature biotechnology 36(5): 421–427.

Hu, Y.; Hase, T.; Li, H. P.; Prabhakar, S.; Kitano, H.; Ng, S. K.; Ghosh, S.; and Wee, L. J. K. 2016. A machine learning approach for the identification of key markers involved in brain development from single-cell transcriptomic data. BMC genomics 17(13): 1025.

Huang, P. H.; Xu, A. M.; and White, F. M. 2009. Oncogenic EGFR signaling networks in glioma. Science signaling 2(87): re6–re6.

Kipf, T. N.; and Welling, M. 2016. Semi-supervised classification with graph convolutional networks. arXiv preprint arXiv:1609.02907.

Langfelder, P.; and Horvath, S. 2008. WGCNA: an R package for weighted correlation network analysis. BMC bioinformatics 9(1): 559.

Liang, W.; Yao, J.; Chen, A.; Lv, Q.; Zanin, M.; Liu, J.; Wong, S.; Li, Y.; Lu, J.; Liang, H.; et al. 2020. Early triage of critically ill COVID-19 patients using deep learning. Nature Communications 11(1): 1–7.

Liaw, A.; Wiener, M.; et al. 2002. Classification and regression by randomForest. R news 2(3): 18–22.

Ngo, T. T.; Moufarrej, M. N.; Rasmussen, M.-L. H.; Camunas-Soler, J.; Pan, W.; Okamoto, J.; Neff, N. F.; Liu, K.; Wong, R. J.; Downes, K.; et al. 2018. Noninvasive blood tests for fetal development predict gestational age and preterm delivery. Science 360(6393): 1133–1136.

Paszke, A.; Gross, S.; Chintala, S.; Chanan, G.; Yang, E.; DeVito, Z.; Lin, Z.; Desmaison, A.; Antiga, L.; and Lerer, A. 2017. Automatic differentiation in pytorch.

Shen, B.; Yi, X.; Sun, Y.; Bi, X.; Du, J.; Zhang, C.; Quan, S.; Zhang, F.; Sun, R.; Qian, L.; et al. 2020. Proteomic and metabolomic characterization of COVID-19 patient sera. Cell.

Srinivas, S.; and Fleuret, F. 2019. Full-gradient representation for neural network visualization. In Advances in Neural Information Processing Systems, 4124–4133.

Suykens, J. A.; and Vandewalle, J. 1999. Least squares support vector machine classifiers. Neural processing letters 9(3): 293–300.

Veličković, P.; Cucurull, G.; Casanova, A.; Romero, A.; Lio, P.; and Bengio, Y. 2017. Graph attention networks. arXiv preprint arXiv:1710.10903.

Wachowiak, R.; Krause, M.; Mayer, S.; Peukert, N.; Suttkus, A.; Müller, W. C.; Lacher, M.; Meixensberger, J.; and Nestler, U. 2018. Increased L1CAM (CD171) levels are associated with glioblastoma and metastatic brain tumors. Medicine 97(38).

Waldrop, C. A.; Youn, S.; and Patterson, L. G. 2014. Network to network interface (NNI) for multiple private network service providers. US Patent 8,756,344.

Wang, T.; Lu, W.; Yang, F.; Liu, L.; Dong, Z.; Tang, W.; Chang, J.; Huan, W.; Huang, K.; and Yao, J. 2020. Microsatellite Instability Prediction of Uterine Corpus Endometrial Carcinoma Based on H&E Histology Whole-Slide Imaging. In 2020 IEEE 17th International Symposium on Biomedical Imaging (ISBI), 1289–1292. IEEE.

Yang, F.; Yi, M.; Liu, Y.; Wang, Q.; Hu, Y.; and Deng, H. 2018. Glutaredoxin-1 silencing induces cell senescence via p53/p21/p16 signaling axis. Journal of proteome research 17(3): 1091–1100.

Yau, T. O. 2019. Precision treatment in colorectal cancer: Now and the future. JGH Open 3(5): 361–369.

Zeng, A.; Hu, Q.; Liu, Y.; Wang, Z.; Cui, X.; Li, R.; Yan, W.; and You, Y. 2015. IDH1/2 mutation status combined with Ki-67 labeling index defines distinct prognostic groups in glioma. Oncotarget 6(30): 30232.

Zhang, H.; Yang, F.; Guo, Y.; Wang, L.; Fang, F.; Wu, H.; Nie, S.; Wang, Y.; Fung, M.-L.; Huang, Y.; et al. 2018. The contribution of chronic intermittent hypoxia to OSAHS: From the perspective of serum extracellular microvesicle proteins. Metabolism 85: 97–108.

Zhao, Y.; Yang, F.; Fang, Y.; Liu, H.; Zhou, N.; Zhang, J.; Sun, J.; Yang, S.; Menze, B.; Fan, X.; et al. 2020. Predicting Lymph Node Metastasis Using Histopathological Images Based on Multiple Instance Learning With Deep Graph Convolution. In Proceedings of the IEEE/CVF Conference on Computer Vision and Pattern Recognition, 4837–4846.

Zhou, Y.; Zhou, B.; Pache, L.; Chang, M.; Khodabakhshi, A. H.; Tanaseichuk, O.; Benner, C.; and Chanda, S. K. 2019. Metascape provides a biologist-oriented resource for the analysis of systems-level datasets. Nature communications 10(1): 1–10.

